# Aspects of flow variability and spatial context predict temporal beta diversity in river metacommunities

**DOI:** 10.1101/2022.09.07.506991

**Authors:** P. Saffarinia, R. Conway, K.E Anderson

## Abstract

1. River catchments are dynamic networks that contain multiple levels of spatial and temporal complexity. Benthic macroinvertebrates are key indicator taxa throughout catchments and beta diversity has been used as a metric to explore determinants of community composition at the catchment scale. Commonly explored drivers of beta diversity include environmental and spatial variables such as flow, temperature, and spatial distance. While factors influencing spatial beta diversity have been explored, factors explaining temporal beta diversity have been understudied. Temporal beta diversity is predicted to also be important since community assembly mechanisms are not stable over time, and more studies are needed to determine which factors most strongly determine temporal beta diversity patterns.
2. We investigated the effects of local environmental variables, flow variability, and spatial context on temporal beta-diversity using a large, publicly available biomonitoring dataset from river networks in California. Data included benthic macroinvertebrate community composition and environmental data from multiple locations and years, allowing us to explore temporal changes in these communities as a function of site-specific environmental and spatial factors. Associated gage data were used to calculate hydrograph metrics and contextualize the flow regime at each location over long timescales. We then used beta regression to model the relationship between benthic macroinvertebrate temporal beta-diversity, environmental variables, flow regimes, and spatial network context.
3. Flow and spatial catchment-related predictors were the strongest predictors of temporal beta diversity, while changes in environmental variables were much weaker. Channel slope, drainage density, and upstream catchment area were the most significant spatial factors. Channel slope showed a negative relationship with temporal beta diversity, while drainage density and upstream catchment area showed positive ones. Temporal beta diversity was also higher when the rate and magnitude of rises and falls in flow was higher in the hydrograph as well as when the number of zero-flow days and the duration of flow rises and falls was higher.
4. Overall, our results indicate that temporal beta diversity of freshwater benthic macroinvertebrates is shaped by both long-term hydrological context and spatial context, and that these factors may serve as better predictors of long term community variability than variability in point estimates of environmental measurements. Flow regimes and spatial metrics may provide more environmental context than point-estimate environmental measurements, as the latter may not accurately capture the dynamic conditions that drive variability in metacommunity responses.
5. Our study supports the need for biomonitoring efforts at long spatial and temporal timescales, and highlights the need to consider metacommunity change in the management of freshwater systems.

## 1 Introduction

Trends in community diversity and distribution are often studied by calculating beta diversity, which can be defined as the dissimilarity in community composition between samples (Anderson et al. 2006). River systems are ideal for the study of the drivers of beta diversity because they exhibit multiple layers of variability across spatial and temporal scales (Tonkin et al. 2018, Harvey and Altermatt 2019). Many classic frameworks in freshwater ecology were attempts to explain spatial and temporal patterns in biodiversity (Vannote and Minshall 1980, Ward and Stanford 1983, Pringle et al. 1988, Townsend 1989) which have been expanded to consider more complex spatial and temporal contexts (Grant et al. 2007, Altermatt 2013, Brown et al. 2015, Green et al. 2022). Despite considerable progress, most beta diversity studies that consider large spatial scales have assumed that community assembly mechanisms are stable over time (Thompson and Townsend 2006, Brown and Swan 2010, Finn et al. 2011, Henriques-Silva et al. 2019). However, stream metacommunities exhibit significant temporal variability in community composition and resulting beta diversity (Datry et al. 2016, Ruhí et al. 2017, Cañedo-Argüelles et al. 2020, Larsen et al. 2021). For example, beta diversity has been shown to vary temporally due to changes in hydrological conditions, which renders assembly mechanisms more stable in perennially flowing streams (Sarremejane et al. 2017b). Additionally, context dependence reported in prior studies could be due to the absence of observations at temporal scales that were long enough to capture overall variability in beta diversity (Tonkin et al. 2016).

The relative influence of temporal processes as drivers of beta diversity continues to be an unresolved issue in ecology; this is especially true in freshwater systems because of their dynamic environmental conditions (Tonkin et al. 2017, Cañedo-Argüelles et al. 2020). Given that flow regimes can affect environmental conditions and community composition in rivers (Power et al. 1995, Poff et al. 1997), spatial and temporal variation in catchment flow regimes can strongly influence beta-diversity patterns (Rolls et al. 2017). Spatial variation in hydrological connectivity, stream discharge magnitude and discharge duration have been shown to be particularly important in the determination of beta-diversity patterns, as these factors not only affect local environmental conditions, but also alter the ability of organisms to disperse throughout river networks (Clarke et al. 2010, Liu et al. 2013, Warfe et al. 2014, Leigh and Datry 2016). For example, differences in the magnitude and duration of peak discharge events driven by channel slope can alter the strength of local environmental filtering (Nippgen et al. 2011). Empirical studies have also shown that freshwater beta diversity is lowest when floods and high flow events are more frequent owing to higher metacommunity connectivity, while community dissimilarity increases when discharge is lost and fragmentation occurs (Bogan et al. 2013, Fazi et al. 2013, Starr et al. 2014, Datry et al. 2014, Cañedo-Argüelles et al. 2020). However, studies of beta-diversity patterns across regions with distinct hydrological characteristics are not found in the literature (Rolls et al. 2017), and they are necessary to better understand the interaction between spatially structured flow regimes and other environmental processes on temporal beta-diversity.

The spatial structure and context of river networks can additionally dictate beta-diversity patterns by driving patterns of environmental variation and influencing dispersal connectivity (Grant et al. 2007, Altermatt 2013, Brown et al. 2017). The metacommunity paradigm, which highlights processes that structure communities beyond the local scale, has facilitated the progress in understanding spatial patterns of beta diversity (Leibold et al. 2004, Winegardner et al. 2012). This is particularly true in freshwater systems, where beta-diversity patterns can strongly respond to spatial structure at local and regional scales (Tonkin et al. 2018). Further, the drivers of benthic macroinvertebrate turnover have been shown to be scale-dependent. Evidence suggests that headwater communities are structured more by local environmental conditions (i.e. high species sorting), while higher-order streams and mainstems are influenced more by distance and corresponding connectivity metrics, such as network position and upstream area (Brown and Swan 2010, Schmera et al. 2017, Brown et al. 2017). However, studies that separate local and regional processes to explain shifts in beta diversity have been unpersuasive; rather, evidence shows that local and regional factors are more connected than originally thought (Heino et al. 2015).

More studies are needed to better understand relationships between temporal beta diversity and the combined effects of changing local environmental conditions, flow regimes, and spatial context. We examined patterns of temporal beta diversity in stream benthic macroinvertebrate communities across multiple years spanning a large geographic area in the Sierra Nevada and Cascade mountain ranges in California, USA. There is a considerable amount of spatial and temporal variation in hydrology and environmental variables throughout this study region, with distinct flow regimes that strongly affect benthic macroinvertebrate assemblages (Lusardi et al. 2016). The importance of different fundamental processes in structuring freshwater communities has been debated, and results may be contingent on the spatial and temporal scale of the study (Heino et al. 2014). Thus, by using a dataset that spans large spatial and temporal scales, we aim to overcome such limitations and improve our understanding of the effects of flow regimes, environmental variation, and spatial context on temporal beta-diversity patterns. Specifically, we asked whether temporal beta diversity 1) reflected changes in local environmental conditions; 2) was influenced by differences in flow regimes and, if so, which aspects weere most important; and 3) was different among sites depending on those sites’ spatial context.

## 2 Methods

### 2.1 Benthic macroinvertebrate and environmental variable data selection

Data on benthic macroinvertebrate community composition and site-level environmental variables were obtained from the California Surface Water Ambient Monitoring Program (SWAMP; https://www.waterboards.ca.gov/water_issues/programs/swamp/), which is publicly accessible through the California Environmental Data Exchange Network (CEDEN, http://www.ceden.org). SWAMP employs a standardized sampling protocol for benthic macroinvertebrates and environmental variables (Ode et al. 2016). These data are largely organized by county. We restricted the data used to counties that generally fall within the Sierra Nevada range and the southern portion of the Cascade mountain range in California, USA, since these regions had the highest number of sites with samples taken over multiple years and a gradient of environmental variation that has previously been shown to influence community composition (Lusardi et al. 2016). The lowest level of taxonomic resolution was genus, though some taxa such as chironomids were only identified to subfamily. Genus is a finer level of taxonomic resolution than used in several prior studies (Heino 2011, Datry et al. 2014, Leigh and Datry 2016). At most SWAMP sampling sites, environmental variables (dissolved oxygen, pH, conductivity, temperature, stream velocity) are measured concurrently with benthic macroinvertebrate sampling.

We mapped all selected sites using ArcGIS (version 9.x) and National Hydrography Dataset (NHDv2) stream layers, taking care to combine site names that were sampled at the same site. To control for seasonal variation in benthic macroinvertebrate metacommunities, selected data were restricted to samples taken between June and September from 1998-2017. Additionally, we only chose sites that had at least two sampling timepoints, though this does not mean the sites were sampled annually. Frequently, sites were sampled once and then resampled after an arbitrary number of years (anywhere between 1-15 years).

Five counties were found to have sufficient spatial and temporal coverage to warrant inclusion in further analyses. While El Dorado, Alpine and Mono counties were all clustered south of Lake Tahoe, CA, Tehama and Plumas counties covered the northern end of the Sierra Nevada range and a small part of the southern Cascade mountain range (Figure 1). The biotic samples had a temporal spread of 1998-2017, with a maximum spatial range of 450km and average total area between all samples of 36,000 km^2^. The dataset included a total of 110 sites across 5 counties (Alpine, El Dorado, Mono, Plumas, and Tehama) with a total of 382 genera, spanning stream orders 1-6.

**Figure 1.**
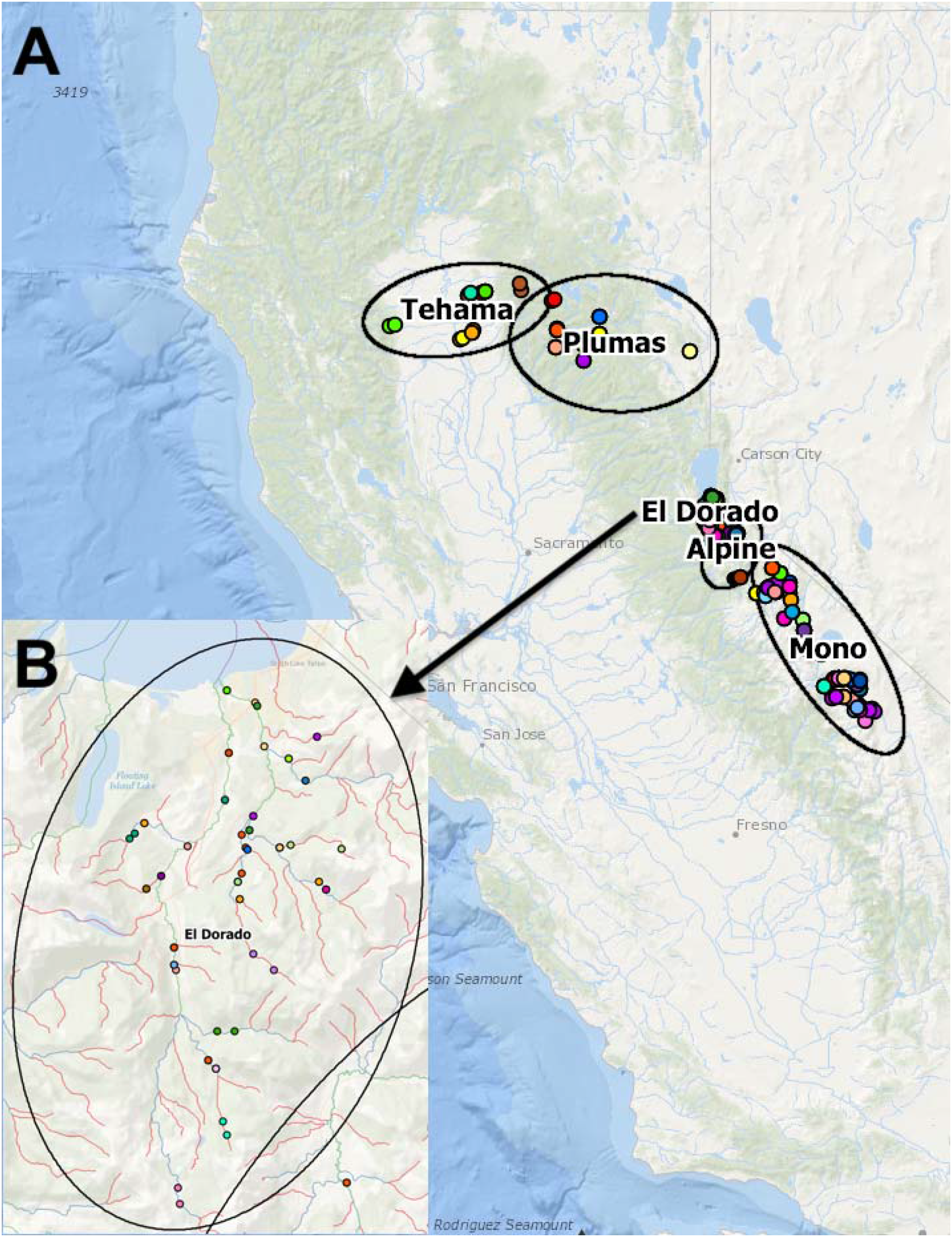
Map of sampling sites used in our analyses. A) Sampling sites were distributed across five counties in the state of California, USA. B) Inset shows an example distribution of sampling sites within a single catchment (El Dorado).

### 2.2 Environmental distance and beta diversity

We calculated environmental and community dissimilarity across sample years at each site. To calculate a metric of environmental distance (environmental dissimilarity), site-specific environmental data were scaled, centered, and ordinated with principal components analysis (PCA) over all timepoints. Principal components with eigenvalues greater than 1 were kept, and environmental distance was defined as the Euclidian distance calculated between pairs of sites in multivariate space (Brown and Swan 2010).

Temporal beta diversity in benthic macroinvertebrate communities was calculated as the Bray-Curtis community dissimilarity of taxa (genus or sub-family) abundances between all timesteps at a given site using the betapart package for R (Baselga 2013).

### 2.3 Analysis of flow regimes

The SWAMP sampling locations were paired with long-term flow data from United States Geological Survey (USGS) stream flow gage sites. Gage locations within a 10 km buffer (Leigh and Datry 2016) and similar stream order to SWAMP sampling sites were identified using the Consortium of Universities for the Advancement of Hydrologic Science (CUAHSI) HydroClient web interface (data.cuahsi.org), and snapping gaged sites to an ArcGIS layer to determine distance to SWAMP sampling locations. Entire hydrograph time series from paired flow gages were used, with an average span of 20 years per gage. The River Analysis Package (RAP 3.0.8) was used to calculate hydrograph metrics and contextualize the flow regime at each location over long timescales (Kennard et al. 2009). Twenty-six flow metrics were calculated (Supplemental Table S1), and then chosen to evaluate critical aspects of flow variability including flow magnitude, duration, frequency, timing and rate of change (Poff et al. 1997, Leigh and Datry 2016). These calculated metrics and their ecological importance have been previously described in detail; see for example (Olden and Poff 2003, Kennard et al. 2009, Leigh and Datry 2016). Metrics calculated from compiled flow data were then centered, scaled, and ordinated using PCA to collapse flow variability information because many different flow metrics were correlated.

### 2.4 Stream catchment metrics

Upstream catchment area from the sampling location, channel slope, drainage density (average length of channel per unit area of catchment) and proportion of land cover types in the upstream catchment area were calculated for each benthic macroinvertebrate sampling location. This was accomplished by constructing a landscape network (LSN) with the functional linkage of water basins and streams (FLoWS) ArcGIS geoprocessing toolset (Theobald et al. 2005), and subsequently, the spatial tools for the analysis of river systems (STARS) toolset was used to generate and format spatial data (Peterson and Ver Hoef 2014, Isaak et al. 2014). Catchment characteristics were derived from NDHPlusv2, while land cover was derived from the National Land Cover Database (NLCD; Supplemental Table S4). Each upstream area was square-root transformed, which is analogous to discharge volume and overall stream size (Leopold and Maddock 1953).

### 2.5 Statistical analyses

The relationships among temporal beta diversity, environmental distance, time between sampling periods (hereafter referred to as Δtime), flow metrics, stream catchment metrics, and land cover were statistically modeled using beta regression. Beta regression was chosen to model community dissimilarity values because they are between the unit interval (0,1) and the data violated assumptions of normality and equal variances (data appeared to be heteroskedastic). Assumptions of normality and equal variance were checked with Shapiro-Wilk’s, Lavene’s test, and bptest in the car package for R (Fox et al. 2022), as well as with Q-Q and Cook’s distance plots. Beta regressions are flexible with respect to heteroskedasticity and incorporate extra precision parameters, which can depend on a set of similar or different regressors to account for extra variance in data (Zeileis and Cribari-Neto 2010). Collinearity between predictors was checked by examining the variance inflation factors (VIFs) in the car package after running beta regressions, and any predictors with a VIF greater than 2 were removed (Fox and Monette 1992). For NLCD land cover data, only %forest cover was used as a predictor because it resulted in the largest model performance improvement, while the inclusion of other land cover data increased the VIF above the acceptable range (Fox and Monette 1992). The addition of a precision parameter in the beta regression model was determined by checking the addition of all predictors as a precision parameter and comparing model performance between models with and without precision parameters with AIC. It was determined that using drainage density as a precision parameter accounted for the most unexplained variance and was used in the final modelling approach.

We used AICc to select the most parsimonious combinations of variables for predicting temporal variation in beta diversity. Using the MuMIn package for R (Barton 2022), we determined which top-ranked models had a ΔAICc less than 4. The relative importance of each predictor was obtained by summed Akaike weights (SW) in the top-ranked models. SW assists in detecting important predictors even if they are not included in the top model when they appear often in other model formulations (Burnham et al. 2010, Giam and Olden 2015). Then, we model-averaged all models with a ΔAICc of less than 4 to account for model selection uncertainty and obtain robust predictions (Grueber et al. 2011), and coefficient estimates from model averaged output were used to understand the importance of predictors. From the averaged beta regression, predicted univariate relationships between each independent variable and beta diversity (mean of the predicted beta distribution) were plotted to visually examine significant effects from the averaged model. The predicted 5% and 95% quantiles were also plotted on these graphs, which revealed increasing precision in the beta regression (i.e., 90% confidence interval around the mean) (Zeileis and Cribari-Neto 2010).

## 3 Results

### 3.1 Ordinations of environmental and flow variables

In the ordinations of sampled environmental variables, the first four principal components explained 88% of the variation, and those were used to calculate Euclidian distance, which was used as the measure of environmental distance among sampled time points in a location (Supplemental Table S2). For the ordination of the flow metrics, the first four principal component axes captured 83% of the variability, so these scores were used in further analyses (Supplemental Table S3). PC1 and PC4 were repeatedly identified as the most important predictors of flow variability (Table 1; Supplemental Table S3). The flow metrics with the strongest loading on PC1 were related to rates and magnitudes of rises and falls in flow over time, while PC4 was related to the number of zero-flow days and the duration of flow rises and falls in the hydrograph. In general, most counties were structured more by PC1, and this axis explained most of the variation in flow metrics (Figure 2). Plumas county sites in particular were structured more by PC1, while Mono county sites were structured more by PC4.

**Figure 2.**
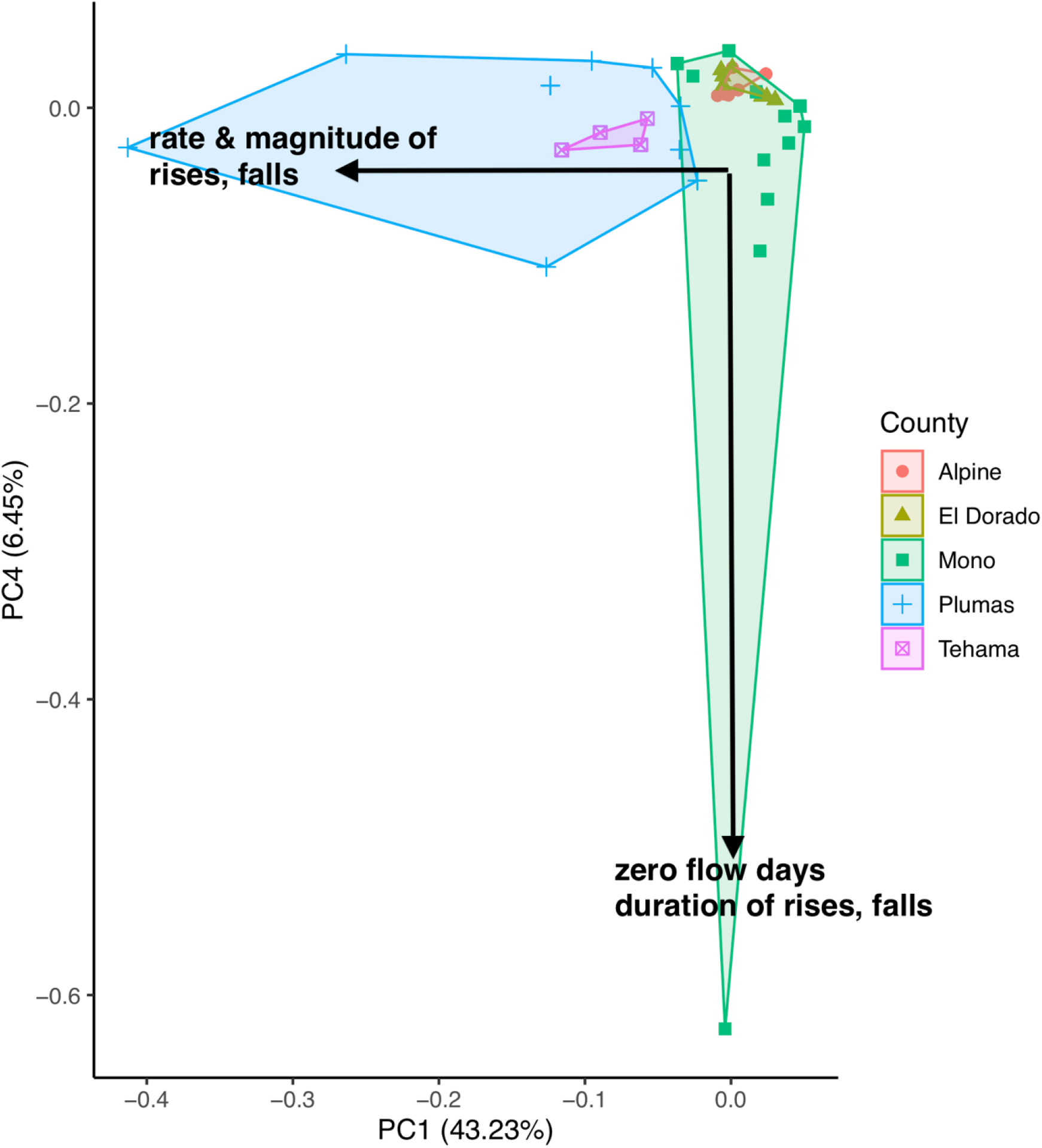
PCA of flow metrics calculated from USGS flow gage data in RAP. PC1 and PC4 are presented since they had the greatest explanatory power in statistical models of beta diversity. Gage locations are grouped by county.

**Table 1.**
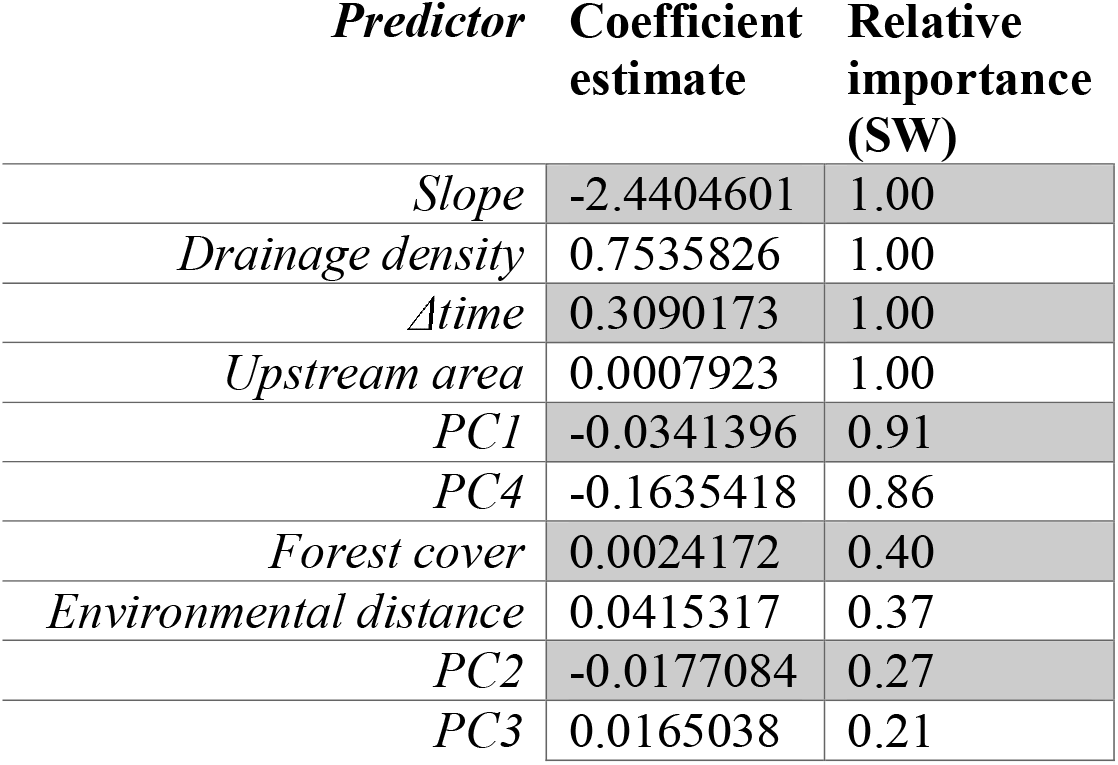
Coefficient estimates from model averaging of beta regressions with a delta AICc of less than 4. The relative importance of each predictor of temporal beta diversity was calculated as how many times that variable was selected in top models.

| *Predictor* | **Coefficient estimate** | **Relative importance (SW)** |
| --- | --- | --- |
| *Slope* | -2.4404601 | 1.00 |
| *Drainage density* | 0.7535826 | 1.00 |
| *Δtime* | 0.3090173 | 1.00 |
| *Upstream area* | 0.0007923 | 1.00 |
| *PC1* | -0.0341396 | 0.91 |
| *PC4* | -0.1635418 | 0.86 |
| *Forest cover* | 0.0024172 | 0.40 |
| *Environmental distance* | 0.0415317 | 0.37 |
| *PC2* | -0.0177084 | 0.27 |
| *PC3* | 0.0165038 | 0.21 |

### 3.2 Beta diversity predictors

Twenty-three beta regression models had ΔAICc < 4 and were therefore selected to explain temporal beta diversity of benthic macroinvertebrates. Out of the suite of the ten selected predictors, six were most important based on summed Akaike weights. Flow and catchment-related predictors had SW values > 0.8, while forest cover, environmental distance, PC2 and PC3 were less important predictors with SW values <0.4 (Table 1).

Drainage density, Δtime, and environmental distance exhibited positive relationships with beta diversity in the model averaged beta regression (estimated coefficients of 0.7536, 0.3090 and 0.0415; Figures 3F, 3A and 3B; Table 1). Upstream area and forest cover exhibited slightly positive relationships with beta diversity (estimated coefficients of 0.0007 and 0.0024; Figure 3E and 3H, Table 1). Channel slope, PC1, and PC4 exhibited negative relationships with beta diversity (estimated coefficients of −2.4405, −0.0341 and −0.1635; Figures 3G, 3C and 3D; Table 1). Since flow metrics were negatively related to PC1 and PC4, the more negative values on the x-axis in Figures 3C and 3D signifies a stronger relationship with that metric. In other words, communities were more dissimilar when the rate and magnitude of rises and falls in flow was higher in the hydrograph, while they were also more dissimilar when the number of zero-flow days and the duration of flow rises and falls was higher.

**Figure 3.**
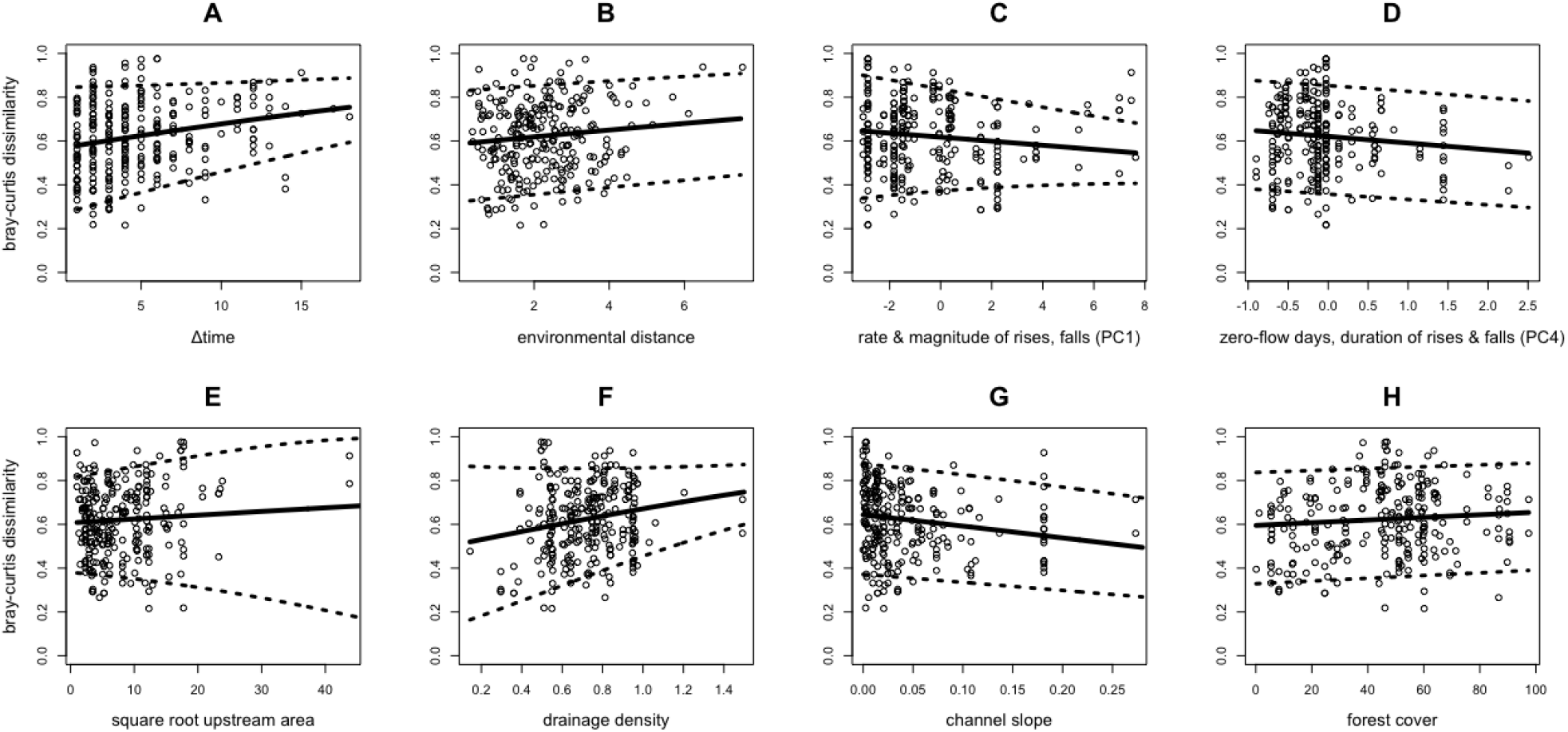
Univariate relationships between selected predictors and temporal beta diversity. Regression lines are derived from the mean of the predicted beta distribution in the averaged model. Dotted lines are 5% and 95% quantiles, showing the 90% confidence intervals.

## 4 Discussion

Although local environmental conditions are known to structure metacommunities in riverine systems (Brown and Swan 2010, Grönroos et al. 2013, Dala-Corte et al. 2017, Gansfort and Traunspurger 2019), point estimates of this historically important predictor were fairly weak in explaining temporal beta diversity in this study. Point estimates in this study included local environmental variables, such as dissolved oxygen and temperature, which were sampled when benthic macroinvertebrates were collected. While both environmental conditions and community composition changed at sites throughout our study, changes in environmental point estimates did not strongly predict beta diversity. Both spatial factors--channel slope, drainage density, upstream catchment area--and long-term flow metrics explained a substantially higher portion of community change. The explanation for this result may be twofold. First, spatial metrics and flow metrics may provide better environmental context by serving as proxies for a more complex suite of variables than point-estimate environmental measurements. For instance, metacommunities inhabiting certain sections of catchments can have unique dispersal abilities that reflects adaptations to different ranges environmental conditions (Bogan et al. 2013). Additionally, point-estimate measurements may not have been able to accurately capture varying local environmental conditions that drive variability in metacommunity responses (Cañedo-Argüelles et al. 2015). This is in contrast to our flow metrics which summarized flow regimes over multiple years.

The two most influential aspects of the hydrograph that predicted higher beta diversity were related to characterizing extreme flow variability at sites: the magnitude and duration of flow rises and falls and the number of zero-flow days (Figures 3C and 3D). It appears that variability in discharge, rather than average discharge at a site, potentially drives more compositional change in these systems. Prior research supports this conclusion, as others have found that metrics involving stream flow loss, such as duration without connected surface flow, act as a strong environmental filter, and drive patterns in benthic macroinvertebrate community composition (Clarke et al. 2010, Stubbington et al. 2019). In other words, when metacommunities are distributed within a catchment that receives high variability in flow patterns, beta diversity increases because communities are repeatedly disturbed. However, it is important to note that the sites within this study primarily exhibited a perennial flow regime, and when metacommunities are more adapted to extreme flow variability (such as in intermittent systems), patterns in beta diversity can be more variable as taxa can exhibit traits to mitigate effects of strong flow variability, such as dispersal strength (Thompson and Townsend 2006, Leigh and Datry 2016).

Local communities in our study exhibited higher temporal beta diversity when occupying areas with higher drainage density and with greater upstream area. By being positioned in more central areas of catchments, these communities could be more dispersal connected and influenced by mass effects (Brown and Swan 2010, Brown et al. 2011, 2017), which could subsequently increase beta diversity. Further, asynchrony between communities could contribute to high compositional change if they occur in areas of high drainage density that facilitates dispersal. With temporally variable local environmental conditions, communities at large spatial scales may fluctuate in diversity through time (Carlson and Satterthwaite 2011, Yeakel et al. 2014), with different patches contributing more or less to beta diversity. Though headwater communities typically experience greater disturbances and more spatial dissimilarity (Finn et al. 2011, Göthe et al. 2013), strong species sorting and limited dispersal connectivity could create a fixed set of taxa that can recolonize such sites (Little and Altermatt 2018). Headwaters tend to be naturally more isolated with lower potential dispersal connectivity (Finn et al. Brown et al. 2011), but those in areas with high drainage density could function similarly to lower mainstem sections that receive input from more patches (Finn et al. 2011, Schmera et al. 2017). Channel slope had a strongly negative relationship with temporal beta diversity, which is consistent with other spatial effects because areas with lower slope are generally associated with mainstems and higher order, lower elevation sites, while sites with a higher slope tend to correspond to higher elevation headwater locations.

Though temporal environmental heterogeneity is known to impact metacommunities (Sarremejane et al. 2017b), we found that time in and of itself, when controlling for temporally structured environment and space, was also a strong predictor of temporal beta diversity in this study. This result could be attributed to imperfect measurement of environmental change and/or lagged responses of different populations. The sampled environmental variables (dissolved oxygen, pH, conductivity, temperature and flow velocity) may not have been the most important environmental drivers of beta diversity in the catchments analyzed in this study (Tolonen et al. 2017). Further, the effects of local environmental changes can take years to move through populations, and the sensitivity and trait profiles of individual taxa are variable through space and over time (Hawkins et al. 2014, Tonkin et al. 2015). Alternatively, the signal of time could be related to ecological drift and neutral processes (Gilbert and Levine 2017). Under more neutral conditions, long-term variation in environmental conditions and morality risk are decoupled from species traits and identity, allowing stochastic changes in community composition to build-up over time, corresponding to increased beta diversity between longer time points. Ecological drift and neutral processes have been found to be important in structuring metacommunities in other freshwater systems (Sarremejane et al. 2017a, Dong et al. 2017, Siqueira et al. 2020).

## 5 Conclusion

Overall, our results indicate that temporal beta diversity of freshwater benthic macroinvertebrates may be strongly shaped by both long-term hydrological context and spatial context, and that these factors may serve as better predictors of long term community variability than variability in point estimates of environmental measurements. Recent research on flow variability has emphasized extreme flow events such as drought and floods as well as intermittency in driving community patterns (Datry et al. 2014, Schneck et al. 2017, Saffarinia et al. 2022, Rolls et al. 2022). In our study, the principal component axis with strong loadings for the magnitude and rates at which the hydrograph rises or falls showed the strongest predictive ability of temporal beta diversity. Thus, not only is it important to focus on extreme flow events, such as zero flow days, but also on the rate at which flow increases and decreases. Further studies are needed to disentangle the importance in the rate of change in flow compared to zero flow days for beta diversity and mechanisms by which these components influence community assembly and change. Our results also point to the importance of conducting these studies to incorporate the temporal variability across whole river networks to properly understand community assembly mechanisms, as these may vary depending on community position and spatial context. Assembly mechanisms can be further elucidated by incorporating changes in functional traits because traits can be directly linked to dispersal and persistence ability in patches of a river network (Boersma et al. 2013). These findings highlight the importance of biomonitoring regimes implemented at long timescales and large spatial scale because discrete point-estimate sampling of local environmental conditions does not always reflect patterns in freshwater metacommunity change (Patrick et al. 2021).

## Acknowledgements

We are grateful to Marko Spasojevic and Jeffery Diez for feedback on study design and statistical analyses, and to Bryan Brown, Eric Sokol, and Chris Swan for inspiring us to think about beta diversity in river networks. We thank Lisa Dong for assistance with data assimilation. The advice of Bill Walton and Matthew Daugherty was helpful in initializing this study.

## Author Contributions

Conceptualization: PS, KEA. Developing methods: PS, KEA, RC. Data analysis: PS, KEA, RC. Preparation of figures and tables: PS. Conducting the research, data interpretation, writing: PS, KEA

## Data Availability Statement

Data are available from the first author (PS) upon reasonable request.

## References

Altermatt, F. 2013. Diversity in riverine metacommunities: a network perspective. Aquatic Ecology 47:365–377.

Anderson, M. J., K. E. Ellingsen, and B. H. McArdle. 2006. Multivariate dispersion as a measure of beta diversity. Ecology Letters 9:683–693.

Barton, K. 2022. Package “mumin.” from CRAN for R.

Baselga, A. 2013. Multiple site dissimilarity quantifies compositional heterogeneity among several sites, while average pairwise dissimilarity may be misleading. Ecography 36:124–128.

Boersma, K. S., M. T. Bogan, B. A. Henrichs, and D. A. Lytle. 2013. Invertebrate assemblages of pools in arid-land streams have high functional redundancy and are resistant to severe drying. Freshwater Biology 59:491–501.

Bogan, M. T., K. S. Boersma, and D. A. Lytle. 2013. Flow intermittency alters longitudinal patterns of invertebrate diversity and assemblage composition in an arid-land stream network. Freshwater Biology 58:1016–1028.

Brown, B. L., and C. M. Swan. 2010. Dendritic network structure constrains metacommunity properties in riverine ecosystems. Journal of Animal Ecology 79:571–580.

Brown, B. L., C. M. Swan, D. A. Auerbach, E. H. C. Grant, N. P. Hitt, K. O. Maloney, and C. Patrick. 2015. Metacommunity theory as a multispecies, multiscale framework for studying the influence of river network structure on riverine communities and ecosystems. Journal of the North American Benthological Society.

Brown, B. L., C. M. Swan, D. A. Auerbach, E. H. Grant, N. P. Hitt, K. O. Maloney, and C. Patrick. 2011. Metacommunity theory as a multispecies, multiscale framework for studying the influence of river network structure on riverine communities and ecosystems. Journal of the North American Benthological Society 30:310–327.

Brown, B. L., C. Wahl, and C. M. Swan. 2017. Experimentally disentangling the influence of dispersal and habitat filtering on benthic invertebrate community structure. Freshwater Biology 63:48–61.

Burnham, K. P., D. R. Anderson, and K. P. Huyvaert. 2010. AIC model selection and multimodel inference in behavioral ecology: some background, observations, and comparisons. Behavioral Ecology and Sociobiology 65:23–35.

Cañedo-Argüelles, M., C. Gutiérrez-Cánovas, R. Acosta, D. Castro-López, N. Cid, P. Fortuño, A. Munné, C. Múrria, A. R. Pimentão, R. Sarremejane, M. Soria, P. Tarrats, I. Verkaik, N. Prat, and N. Bonada. 2020. As time goes by: 20 years of changes in the aquatic macroinvertebrate metacommunity of Mediterranean river networks. Journal of Biogeography 47:1861–1874.

Cañedo-Argüelles, M., K. S. Boersma, M. T. Bogan, J. D. Olden, I. Phillipsen, T. A. Schriever, and D. A. Lytle. 2015. Dispersal strength determines meta-community structure in a dendritic riverine network. Journal of Biogeography 42:778–790.

Carlson, S. M., and W. H. Satterthwaite. 2011. Weakened portfolio effect in a collapsed salmon population complex. Canadian Journal of Fisheries and Aquatic Sciences 68:1579–1589.

Clarke, A., R. Mac Nally, N. Bond, and P. S. Lake. 2010. Flow permanence affects aquatic macroinvertebrate diversity and community structure in three headwater streams in a forested catchment. Canadian Journal of Fisheries and Aquatic Sciences 67:1649–1657.

Dala-Corte, R. B., F. G. Becker, and A. S. Melo. 2017. The importance of metacommunity processes for long-term turnover of riffle-dwelling fish assemblages depends on spatial position within a dendritic network. Canadian Journal of Fisheries and Aquatic Sciences 74:101–115.

Datry, T., A. S. Melo, N. Moya, J. Zubieta, E. De la Barra, and T. Oberdorff. 2016. Metacommunity patterns across three Neotropical catchments with varying environmental harshness. Freshwater Biology 61:277–292.

Datry, T., S. T. Larned, K. M. Fritz, M. T. Bogan, P. J. Wood, E. I. Meyer, and A. N. Santos. 2014. Broad-scale patterns of invertebrate richness and community composition in temporary rivers: effects of flow intermittence. Ecography 37:94–104.

Dong, X., D. A. Lytle, J. D. Olden, T. A. Schriever, and R. Muneepeerakul. 2017. Importance of neutral processes varies in time and space: Evidence from dryland stream ecosystems. 12:e0176949–14.

Fazi, S., E. Vázquez, E. O. Casamayor, S. Amalfitano, and A. Butturini. 2013. Stream Hydrological Fragmentation Drives Bacterioplankton Community Composition. 8:e64109–10.

Finn, D. S., N. Bonada, C. Múrria, and J. M. Hughes. 2011. Small but mighty: headwaters are vital to stream network biodiversity at two levels of organization. Journal of the North American Benthological Society 30:963–980.

Fox, J., and G. Monette. 1992. Generalized Collinearity Diagnostics. Journal of the American Statistical Association 87:178–183.

Fox, J., G. G. Friendly, and S. Graves. 2022. The car package. From CRAN for R.

Gansfort, B., and W. Traunspurger. 2019. Environmental factors and river network position allow prediction of benthic community assemblies: A model of nematode metacommunities. Nature Publishing Group 9:14716–10.

Giam, X., and J. D. Olden. 2015. Quantifying variable importance in a multimodel inference framework. Methods in Ecology and Evolution 7:388–397.

Gilbert, B., and J. M. Levine. 2017. Ecological drift and the distribution of species diversity. Proceedings of the Royal Society B: Biological Sciences 284:20170507–10.

Göthe, E., N. Friberg, M. Kahlert, J. Temnerud, and L. Sandin. 2013. Headwater biodiversity among different levels of stream habitat hierarchy. Biodiversity and Conservation 23:63–80.

Grant, E. H. C., W. H. Lowe, and W. F. Fagan. 2007. Living in the branches: population dynamics and ecological processes in dendritic networks. Ecology Letters 10:165–175.

Green, M. D., K. E. Anderson, D. B. Herbst, and M. J. Spasojevic. 2022. Rethinking biodiversity patterns and processes in stream ecosystems. Ecological Monographs 92:e1520.

Grönroos, M., J. Heino, T. Siqueira, V. L. Landeiro, J. Kotanen, and L. M. Bini. 2013. Metacommunity structuring in stream networks: roles of dispersal mode, distance type, and regional environmental context. Ecology and Evolution 3:4473–4487.

Grueber, C. E., S. Nakagawa, R. J. Laws, and I. G. Jamieson. 2011. Multimodel inference in ecology and evolution: challenges and solutions. Journal of Evolutionary Biology 24:699–711.

Harvey, E., and F. Altermatt. 2019. Regulation of the functional structure of aquatic communities across spatial scales in a major river network. Ecology 100:e02633–9.

Hawkins, C. P., H. Mykrä, J. Oksanen, and J. J. Vander Laan. 2014. Environmental disturbance can increase beta diversity of stream macroinvertebrate assemblages. Global Ecology and Biogeography 24:483–494.

Heino, J. 2011. A macroecological perspective of diversity patterns in the freshwater realm. Freshwater Biology 56:1703–1722.

Heino, J., A. S. Melo, and L. M. Bini. 2014. Reconceptualising the beta diversity-environmental heterogeneity relationship in running water systems. Freshwater Biology 60:223–235.

Heino, J., A. S. Melo, L. M. Bini, F. Altermatt, S. A. Al-Shami, D. G. Angeler, N. Bonada, C. Brand, M. Callisto, K. Cottenie, O. Dangles, D. Dudgeon, A. Encalada, E. Göthe, M. Grönroos, N. Hamada, D. Jacobsen, V. L. Landeiro, R. Ligeiro, R. T. Martins, M. L. Miserendino, C. S. Md Rawi, M. E. Rodrigues, F. de O. Roque, L. Sandin, D. Schmera, L. F. Sgarbi, J. P. Simaika, T. Siqueira, R. M. Thompson, and C. R. Townsend. 2015. A comparative analysis reveals weak relationships between ecological factors and beta diversity of stream insect metacommunities at two spatial levels. Ecology and Evolution 5:1235–1248.

Henriques-Silva, R., M. Logez, N. Reynaud, P. A. Tedesco, S. Brosse, S. R. Januchowski-Hartley, T. Oberdorff, and C. Argillier. 2019. A comprehensive examination of the network position hypothesis across multiple river metacommunities. Ecography 42:284–294.

Isaak, D. J., E. E. Peterson, J. M. Ver Hoef, S. J. Wenger, J. A. Falke, C. E. Torgersen, C. Sowder, E. A. Steel, M.-J. Fortin, C. E. Jordan, A. S. Ruesch, N. Som, and P. Monestiez. 2014. Applications of spatial statistical network models to stream data. Wiley Interdisciplinary Reviews: Water 1:277–294.

Kennard, M. J., S. J. Mackay, B. J. Pusey, J. D. Olden, and N. Marsh. 2009. Quantifying uncertainty in estimation of hydrologic metrics for ecohydrological studies. River Research and Applications 16:n/a–n/a.

Larsen, S., L. Comte, A. F. Filipe, M.-J. Fortin, C. Jacquet, R. Ryser, P. A. Tedesco, U. Brose, T. Eros, X. Giam, K. Irving, A. Ruhí, S. Sharma, and J. D. Olden. 2021. The geography of metapopulation synchrony in dendritic river networks. Ecology Letters 24:791–801.

Leibold, M. A., M. Holyoak, N. Mouquet, P. Amarasekare, J. M. Chase, M. F. Hoopes, R. D. Holt, J. B. Shurin, R. Law, D. Tilman, M. Loreau, and A. Gonzalez. 2004. The metacommunity concept: a framework for multi-scale community ecology. Ecology Letters 7:601–613.

Leigh, C., and T. Datry. 2016. Drying as a primary hydrological determinant of biodiversity in river systems: a broad-scale analysis. Ecography 40:487–499.

Leopold, L. B., and T. Maddock. 1953. The hydraulic geometry of stream channels and some physiographic implications.

Little, C. J., and F. Altermatt. 2018. Do priority effects outweigh environmental filtering in a guild of dominant freshwater macroinvertebrates? Proceedings. Biological sciences 285:20180205–9.

Liu, J., J. Soininen, B.-P. Han, and S. A. J. Declerck. 2013. Effects of connectivity, dispersal directionality and functional traits on the metacommunity structure of river benthic diatoms. Journal of Biogeography 40:2238–2248.

Lusardi, R. A., M. T. Bogan, P. B. Moyle, and R. A. Dahlgren. 2016. Environment shapes invertebrate assemblage structure differences between volcanic spring-fed and runoff rivers in northern California. Freshwater Science 35:1010–1022.

Nippgen, F., B. L. McGlynn, L. A. Marshall, and R. E. Emanuel. 2011. Landscape structure and climate influences on hydrologic response. Water Resources Research 47:164–17.

Ode, P. R., A. E. Fetscher, and L. B. Buusse. 2016. Standard Operating Procedures (SOP) for the Collection of Field Data for Bioassessments of California Wadeable Streams: Benthic Macroinvertebrates, Algae, and Physical Habitat.

Olden, J. D., and N. L. Poff. 2003. Redundancy and the choice of hydrologic indices for characterizing streamflow regimes. River Research and Applications 19:101–121.

Patrick, C. J., K. E. Anderson, B. L. Brown, C. P. Hawkins, A. Metcalfe, P. Saffarinia, T. Siqueira, C. M. Swan, J. D. Tonkin, and L. L. Yuan. 2021. The application of metacommunity theory to the management of riverine ecosystems. Wiley Interdisciplinary Reviews: Water 8:e1557.

Peterson, E., and J. M. Ver Hoef. 2014. STARS: An ArcGIS toolset used to calculate the spatial information needed to fit spatial statistical models to stream network data. Journal of Statistical Software 56.

Poff, N. L., J. D. Allan, M. B. Bain, J. R. Karr, K. L. Prestegaard, B. D. Richter, R. E. Sparks, and J. C. Stromberg. 1997. The natural flow regime. BioScience 47:769–784.

Power, M. E., A. Sun, G. Parker, W. E. Dietrich, and J. T. Wootton. 1995. Hydraulic Food-Chain Models. BioScience 45:159–167.

Pringle, C. M., R. J. Naiman, G. Bretschko, J. R. Karr, M. W. Oswood, J. R. Webster, R. L. Welcomme, and M. J. Winterbourn. 1988. Patch Dynamics in Lotic Systems - the Stream as a Mosaic. Journal of the North American Benthological Society 7:503–524.

Rolls, R. J., B. C. Chess am, J. Heino, B. Wolfenden, L. A. Thurtell, K. J. M. Cheshire, D. Ryan, G. Butler, I. Growns, and G. Curwen. 2022. Change in beta diversity of riverine fish during and after supra-seasonal drought. Landscape Ecology 37:1633–1651.

Rolls, R. J., J. Heino, D. S. Ryder, B. C. Chess am, I. O. Growns, R. M. Thompson, and K. B. Gido. 2017. Scaling biodiversity responses to hydrological regimes. Biological Reviews 93:971–995.

Ruhí, A., T. Datry, and J. L. Sabo. 2017. Interpreting beta-diversity components over time to conserve metacommunities in highly dynamic ecosystems. Conservation Biology 31:1459–1468.

Saffarinia, P., K. E. Anderson, and D. B. Herbst. 2022. Effects of experimental multi-season drought on abundance, richness, and beta diversity patterns in perennially flowing stream insect communities. Hydrobiologia 849:879–897.

Sarremejane, R., H. Mykrä, N. Bonada, J. Aroviita, and T. Muotka. 2017a. Habitat connectivity and dispersal ability drive the assembly mechanisms of macroinvertebrate communities in river networks. Freshwater Biology 62:1073–1082.

Sarremejane, R., M. Cañedo-Argüelles, N. Prat, H. Mykrä, T. Muotka, and N. Bonada. 2017b. Do metacommunities vary through time? Intermittent rivers as model systems. Journal of Biogeography 44:2752–2763.

Schmera, D., D. Árva, P. Boda, E. Bódis, Á. Bolgovics, G. Borics, A. Csercsa, C. Deák, E. Á. Krasznai, B. A. Lukács, P. Mauchart, A. Móra, P. Sály, A. Specziár, K. Süveges, I. Szivák, P. Takács, M. Tóth, G. Várbíró, A. E. Vojtkó, and T. Erős. 2017. Does isolation influence the relative role of environmental and dispersal-related processes in stream networks? An empirical test of the network position hypothesis using multiple taxa. Freshwater Biology 63:74–85.

Schneck, F., K. Lange, A. S. Melo, C. R. Townsend, and C. D. Matthaei. 2017. Effects of a natural flood disturbance on species richness and beta diversity of stream benthic diatom communities. Aquatic Ecology 51:557–569.

Siqueira, T., V. S. Saito, L. M. Bini, A. S. Melo, D. K. Petsch, V. L. Landeiro, K. T. Tolonen, J. Jyrkänkallio-Mikkola, J. Soininen, and J. Heino. 2020. Community size can affect the signals of ecological drift and niche selection on biodiversity. Ecology 101:e03014.

Starr, S. M., J. P. Benstead, and R. A. Sponseller. 2014. Spatial and temporal organization of macroinvertebrate assemblages in a lowland floodplain ecosystem. Landscape Ecology 29:1017–1031.

Stubbington, R., R. Sarremejane, and T. Datry. 2019. Alpha and beta diversity of connected benthic–subsurface invertebrate communities respond to drying in dynamic river ecosystems. Ecography 42:2060–2073.

Theobald, D., J. Norman, E. Peterson, and S. Ferraz. 2005. Functional linkage of watersheds and streams (FLoWS): network-based ArcGIS tools to analyze freshwater ecosystems. Fisheries.

Thompson, R., and C. Townsend. 2006. A truce with neutral theory: local deterministic factors, species traits and dispersal limitation together determine patterns of diversity in stream invertebrates. Journal of Animal Ecology 75:476–484.

Tolonen, K. E., K. Leinonen, J. Erkinaro, and J. Heino. 2017. Ecological uniqueness of macroinvertebrate communities in high-latitude streams is a consequence of deterministic environmental filtering processes. Aquatic Ecology 52:17–33.

Tonkin, J. D., A. Sundermann, S. C. Jähnig, and P. Haase. 2015. Environmental Controls on River Assemblages at the Regional Scale: An Application of the Elements of Metacommunity Structure Framework. 10:e0135450–19.

Tonkin, J. D., D. M. Merritt, J. D. Olden, L. V. Reynolds, and D. A. Lytle. 2018. Flow regime alteration degrades ecological networks in riparian ecosystems. Nature Ecology & Evolution 1–10.

Tonkin, J. D., J. Heino, A. Sundermann, P. Haase, and S. C. Jähnig. 2016. Context dependency in biodiversity patterns of central German stream metacommunities. Freshwater Biology 61:607–620.

Tonkin, J. D., J. Heino, and F. Altermatt. 2017. Metacommunities in river networks: The importance of network structure and connectivity on patterns and processes. Freshwater Biology 63:1–5.

Townsend, C. R. 1989. The Patch Dynamics Concept of Stream Community Ecology. Journal of the North American Benthological Society 8:36–50.

Vannote, R. L., and G. W. Minshall. 1980. The river continuum concept. Canadian journal of Fisheries and Aquatic Sciences 37:130–137.

Ward, J. V., and J. A. Stanford. 1983. The serial discontinuity concept of lotic ecosystems. Pages 29–42 in T. D. Fontaine and S. M. Bartell (editors). Ann Arbor Scientific Publishers.

Warfe, D. M., S. A. Hardie, A. R. Uytendaal, C. J. Bobbi, and L. A. Barmuta. 2014. The ecology of rivers with contrasting flow regimes: identifying indicators for setting environmental flows. Freshwater Biology 59:2064–2080.

Winegardner, A. K., B. K. Jones, I. S. Y. Ng, T. Siqueira, and K. Cottenie. 2012. The terminology of metacommunity ecology. Trends in Ecology & Evolution 27:253–254.

Yeakel, J. D., J. W. Moore, P. R. Guimarães, and M. A. M. de Aguiar. 2014. Synchronisation and stability in river metapopulation networks. Ecology Letters 17:273–283.

Zeileis, A., and F. Cribari-Neto. 2010. Beta regression in R. Journal of Statistical Software.

