## Supplement for "Aspects of flow variability and spatial context predict temporal beta diversity in river metacommunities"

Supplementary information in support of: Saffarinia, P., R. Conway and Anderson, K.E. Aspects of flow variability and spatial context predict temporal beta diversity in river metacommunities. Submission to Freshwater Biology.

**Supplementary Tables**

**Table S1.** List of all hydrograph metrics calculated from the RAP program (3.0.8). ARI represents annual flood return interval.

Flow metrics calculated with RAP

Minimum

Maximum

Zeros

Longest High Spell

Mean of High Spell Peaks

Mean Duration of High Spell

Mean period Between High Spells

Longest Low Spell

Mean of Low Spell troughs

Mean Duration of Low Spell

Mean period Between Low Spells

Mean magnitude of Rises

Mean duration of Rises

Mean rate of Rise

Mean magnitude of Falls

Mean duration of Falls

Mean rate of Fall

Baseflow Index

Flood Flow Index

Mean Daily Baseflow

Predictability based on monthly mean daily flow

Constancy based on monthly mean daily flow

Contingency based on monthly mean daily flow

Partial series 1 Yr ARI

Partial series 2 Yr ARI

Partial series 10 Yr ARI

**Table S2.** Principal component loadings for environmental variables used followed with the proportion of variance explained by each principal component in the last row.

|  | PC1 | PC2 | PC3 | PC4 | PC5 |
| --- | --- | --- | --- | --- | --- |
| DO | 0.22918951 | -0.6192139 | -0.5660306 | 0.45937268 | -0.1806443 |
| pH | -0.3527708 | -0.3756927 | -0.3639793 | -0.7757993 | 0.00788929 |
| Conductivity | -0.6308911 | -0.0238682 | -0.1406844 | 0.37121591 | 0.66619324 |
| Temperature | -0.6512056 | -0.023585 | 0.16739707 | 0.22182248 | -0.7057961 |
| Velocity | 0.03054022 | -0.6886967 | 0.70662195 | -0.0102799 | 0.1591976 |
| **Proportion of variance explained** | 0.29 | 0.21 | 0.2 | 0.18 | 0.13 |

**Table S3.** Principal component loadings for calculated flow metrics used, followed with the proportion of variance explained by each principal component in the last row. Principal components with a .01 and lower value were omitted.

|  | PC1 | PC2 | PC3 | PC4 | PC5 | PC6 | PC7 | PC8 |
| --- | --- | --- | --- | --- | --- | --- | --- | --- |
| Minimum | 0.10763571 | -0.3067269 | 0.12178947 | 0.0055972 | 0.14277948 | 0.15534737 | 0.06880811 | 0.18050554 |
| Maximum | 0.26129122 | -0.0658395 | -0.1240284 | -0.0552132 | -0.1108631 | 0.11665949 | 0.35030272 | 0.18773578 |
| Zeros | 0.01817507 | 0.04902091 | 0.12161973 | 0.69195636 | -0.0434547 | -0.3413057 | 0.14185927 | -0.1182954 |
| Longest.High.Spell | 0.03913076 | 0.28682591 | -0.1898823 | -0.0005434 | -0.4837625 | 0.06190596 | 0.02731124 | -0.0486084 |
| Mean.of.High.Spell.Peaks | 0.27070362 | -0.0779048 | -0.0802335 | 0.00863112 | -0.0983561 | -0.112447 | 0.15780414 | -0.2170927 |
| Mean.Duration.of.High.Spell | -0.0530403 | 0.26274533 | -0.2025284 | 0.07437964 | -0.3683816 | 0.45899102 | -0.065448 | -0.3399976 |
| Mean.period.Between.High.Spells | -0.1194137 | -0.3055895 | 0.16259944 | 0.00055261 | 0.04430429 | -0.0602718 | 0.38625483 | -0.0291808 |
| Longest.Low.Spell | 0.1980528 | 0.1994047 | -0.1838772 | -0.0356627 | -0.1410055 | -0.1341739 | 0.21871148 | 0.4119322 |
| Mean.of.Low.Spell.troughs | 0.26101769 | -0.1224503 | -0.0922329 | -0.0252612 | 0.035385 | -0.0283359 | 0.18807373 | -0.3052 |
| Mean.Duration.of.Low.Spell | 0.09925961 | 0.2632288 | -0.122849 | -0.0604098 | 0.38288373 | 0.0685444 | 0.3880618 | -0.2035917 |
| Mean.period.Between.Low.Spells | -0.0935771 | -0.0873144 | -0.5670247 | 0.00624925 | 0.30152888 | -0.0461863 | 0.02915868 | -0.0849658 |
| Mean.magnitude.of.Rises | 0.28365368 | -0.0971741 | -0.0255664 | 0.03626232 | 0.00030175 | 0.05728457 | -0.1373915 | 0.00271245 |
| Mean.duration.of.Rises | -0.0602464 | 0.00146731 | -0.5938644 | 0.10444421 | 0.23687225 | -0.1410236 | -0.2361274 | 0.16235909 |
| Mean.rate.of.Rise | 0.26971061 | -0.1084571 | 0.03372238 | 0.07105046 | 0.01124517 | 0.00529296 | -0.3571132 | 0.07083916 |
| Mean.magnitude.of.Falls | 0.28364212 | -0.0971614 | -0.0257176 | 0.03628754 | 3.83E-05 | 0.05715243 | -0.136915 | 0.003005 |
| Mean.duration.of.Falls | 0.02599091 | 0.1073397 | -0.0184499 | 0.66966692 | 0.1324006 | 0.30172336 | 0.05238293 | 0.19157916 |
| Mean.rate.of.Fall | 0.2654238 | -0.0944195 | 0.03480445 | 0.07166278 | -0.0125829 | -0.1223987 | -0.3622718 | -0.1635698 |
| Baseflow.Index | -0.1458783 | -0.3095278 | -0.1883356 | -0.0054907 | -0.1505132 | 0.00476063 | -0.013592 | 0.01099444 |
| Flood.Flow.Index | 0.14587834 | 0.30952783 | 0.18833555 | 0.00549071 | 0.15051316 | -0.0047606 | 0.01359197 | -0.0109944 |
| Mean.Daily.Baseflow | 0.24978406 | -0.1267431 | -0.0806735 | 0.02075437 | -0.0372012 | -0.2463986 | 0.05088907 | -0.4748028 |
| Predictability.based.on.monthly.mean.daily.flow | -0.1507747 | -0.2745413 | -0.0048402 | 0.0970745 | 0.08876926 | 0.44658262 | 0.01999314 | -0.1898235 |
| Constancy.based.on.monthly.mean.daily.flow | -0.157503 | -0.3043942 | -0.060836 | 0.11413825 | -0.1228338 | 0.18655379 | 0.05148722 | -0.0648058 |
| Contingency.based.on.monthly.mean.daily.flow | 0.11769925 | 0.25349299 | 0.13077886 | -0.1043837 | 0.41573674 | 0.2848474 | -0.0862618 | -0.1521959 |
| Partial.series.1.Yr.ARI | 0.27313307 | -0.098082 | 0.03746303 | 0.00704827 | 0.02929325 | 0.15714565 | -0.1506529 | 0.10405893 |
| Partial.series.2.Yr.ARI | 0.27999832 | -0.0947389 | -0.0368289 | -0.0115735 | -0.0317143 | 0.16964212 | 0.03982038 | 0.13393122 |
| Partial.series.10.Yr.ARI | 0.27240429 | -0.0842976 | -0.0916633 | -0.0266708 | -0.084429 | 0.16510515 | 0.2145474 | 0.1674489 |
| Proportion of variance explained | 0.43 | 0.26 | 0.08 | 0.06 | 0.05 | 0.03 | 0.02 | 0.02 |

**Table S4.** Output of dredge function from MuMIn package for R from the full beta regression. All combinations of predictors were included and removed, while only models with a delta AICc of less than 4 are presented. Cells with a value of 0 represent a predictor that was dropped from that model.

| intercept | Δtime | drainage density | environmental distance | forest cover | PC1 | PC2 | PC3 | PC4 | slope | upstream area | df | Loglik | AICc | delta AICc | weight |
| --- | --- | --- | --- | --- | --- | --- | --- | --- | --- | --- | --- | --- | --- | --- | --- |
| -0.241 | 0.04 | 0.754 | 0 | 0 | -0.035 | 0 | 0 | -0.164 | -2.411 | 0.001 | 9 | 133.273 | -247.82 | 0 | 0.082 |
| -0.289 | 0.04 | 0.674 | 0 | 0.002 | -0.034 | 0 | 0 | -0.16 | -2.5 | 0.001 | 10 | 134.209 | -247.528 | 0.293 | 0.071 |
| -0.35 | 0.039 | 0.777 | 0.042 | 0 | -0.035 | 0 | 0 | -0.158 | -2.426 | 0.001 | 10 | 133.966 | -247.041 | 0.78 | 0.056 |
| -0.277 | 0.039 | 0.815 | 0 | 0 | -0.032 | -0.019 | 0 | -0.179 | -2.43 | 0.001 | 10 | 133.843 | -246.795 | 1.026 | 0.049 |
| -0.399 | 0.04 | 0.7 | 0.042 | 0.002 | -0.034 | 0 | 0 | -0.154 | -2.507 | 0.001 | 11 | 134.906 | -246.738 | 1.082 | 0.048 |
| -0.236 | 0.04 | 0.743 | 0 | 0 | -0.035 | 0 | 0.021 | -0.152 | -2.322 | 0.001 | 10 | 133.49 | -246.09 | 1.731 | 0.035 |
| -0.311 | 0.04 | 0.731 | 0 | 0.002 | -0.033 | -0.015 | 0 | -0.172 | -2.499 | 0.001 | 11 | 134.528 | -245.983 | 1.838 | 0.033 |
| -0.376 | 0.038 | 0.833 | 0.039 | 0 | -0.032 | -0.018 | 0 | -0.172 | -2.436 | 0.001 | 11 | 134.436 | -245.798 | 2.022 | 0.03 |
| -0.285 | 0.04 | 0.675 | 0 | 0.002 | -0.034 | 0 | 0.007 | -0.157 | -2.466 | 0.001 | 11 | 134.229 | -245.385 | 2.435 | 0.024 |
| -0.342 | 0.039 | 0.767 | 0.04 | 0 | -0.035 | 0 | 0.018 | -0.148 | -2.346 | 0.001 | 11 | 134.126 | -245.178 | 2.642 | 0.022 |
| -0.287 | 0.037 | 0.869 | 0 | 0 | -0.036 | 0 | 0 | 0 | -2.624 | 0.001 | 8 | 130.866 | -245.153 | 2.667 | 0.022 |
| -0.342 | 0.038 | 0.788 | 0 | 0.003 | -0.036 | 0 | 0 | 0 | -2.72 | 0.001 | 9 | 131.899 | -245.072 | 2.749 | 0.021 |
| -0.413 | 0.039 | 0.749 | 0.039 | 0.002 | -0.033 | -0.013 | 0 | -0.164 | -2.502 | 0.001 | 12 | 135.146 | -245.018 | 2.802 | 0.02 |
| -0.272 | 0.039 | 0.803 | 0 | 0 | -0.032 | -0.018 | 0.018 | -0.168 | -2.352 | 0.001 | 11 | 133.999 | -244.925 | 2.896 | 0.019 |
| -0.18 | 0.042 | 0.677 | 0 | 0 | 0 | 0 | 0 | -0.172 | -2.122 | 0.001 | 8 | 130.692 | -244.806 | 3.015 | 0.018 |
| -0.406 | 0.036 | 0.891 | 0.047 | 0 | -0.036 | 0 | 0 | 0 | -2.624 | 0.001 | 9 | 131.716 | -244.706 | 3.114 | 0.017 |
| -0.463 | 0.037 | 0.81 | 0.047 | 0.003 | -0.036 | 0 | 0 | 0 | -2.708 | 0.001 | 10 | 132.769 | -244.647 | 3.173 | 0.017 |
| -0.396 | 0.04 | 0.7 | 0.041 | 0.002 | -0.034 | 0 | 0.003 | -0.152 | -2.49 | 0.001 | 12 | 134.91 | -244.547 | 3.273 | 0.016 |
| -0.227 | 0.043 | 0.596 | 0 | 0.002 | 0 | 0 | 0 | -0.168 | -2.215 | 0.001 | 9 | 131.613 | -244.5 | 3.32 | 0.016 |
| -0.23 | 0.041 | 0.761 | 0 | 0 | 0 | -0.025 | 0 | -0.189 | -2.177 | 0.001 | 9 | 131.605 | -244.484 | 3.336 | 0.016 |
| -0.276 | 0.037 | 0.839 | 0 | 0 | -0.036 | 0 | 0.036 | 0 | -2.447 | 0.001 | 9 | 131.533 | -244.339 | 3.481 | 0.014 |
| -0.29 | 0.041 | 0.701 | 0.042 | 0 | 0 | 0 | 0 | -0.165 | -2.15 | 0.001 | 9 | 131.38 | -244.034 | 3.787 | 0.012 |
| -0.368 | 0.038 | 0.823 | 0.037 | 0 | -0.033 | -0.017 | 0.015 | -0.163 | -2.367 | 0.001 | 12 | 134.55 | -243.827 | 3.993 | 0.011 |

**Table S4.** Variables from NLCD tested in beta regression.

| **NLCD Value** | **Corresponding habitat type** |
| --- | --- |
| NLCD21PC | Developed, open space |
| NLCD31PC | Barren land |
| NLCD42PC | Evergreen forest |
| NLCD52PC | Shrub/scrub |
| NLCD71PC | Grassland/herbaceous |
